# Neural mechanisms for visuomotor co-regulation in social synchronization

**DOI:** 10.1101/2025.05.21.655276

**Authors:** Stefano Uccelli, Lucia Maria Sacheli, Claudia De Bernardi, Francantonio Devoto, Gianpaolo Basso, Eraldo Paulesu

**Affiliations:** Psychology Department and Milan Center for Neuroscience, University of Milano-Bicocca, Milano, Italy; IRCCS Orthopedic Institute Galeazzi, Milan, Italy; School of Medicine and Surgery, University of Milano-Bicocca, Milano, Italy; Neuroradiology Unit, Fondazione IRCCS San Gerardo dei Tintori, Monza, Italy

**Author notes:** corresponding authors: Stefano Uccelli - Eraldo Paulesu - Department of Psychology, University of Milano-Bicocca P.za Dell’Ateneo Nuovo 1, 20126, Milan (IT).

**Keywords:** entrainment, finger-tapping, interpersonal synchronization, joint action, visuomotor control

## Abstract

During social activities, people coordinate their movements by exchanging visuomotor information. Interpersonal coordination occurs from two complementary processes entailing voluntary (planned) and spontaneous (emergent) synchronization, of which neurofunctional underpinnings are unknown. We investigated the brain correlates of these two synchronization processes during an fMRI finger-tapping task. Dyads formed by IN and OUT scanner participants were instructed to reproduce a target tempo and concurrently try to synchronize to (Joint Action, JA) or resist synchronization with (Non-interactive, NI) the partner’s tapping, whose hand was always visible to the IN participant. Faster, slower, or equal tempi were used to induce within-dyad co-regulation. Results revealed the emergence of tempo contagion: participants tapped faster in response to the partner’s faster taps and vice versa for slower taps. The magnitude of such an interpersonal contagion effect was similar across conditions but associated with the activity of different neural structures. Tempo contagion correlated positively with lateral occipitotemporal cortex (LOTc) activations in the JA condition but negatively with cerebellar activations in the NI condition. This suggests visuomotor information is exploited in opposite ways depending on the task instructions: the LOTc promotes co-regulation to achieve synchronization in JA, whereas the cerebellum prevents tempo contagion to preserve individual stability in NI. This latter result is supported by a negative functional connectivity between the cerebellum and LOTc. These findings have implications for understating the interplay between planned and emergent synchronization during motor interactions.

## Introduction

Synchronizing movements is fundamental during social interactions. Depending on the sought outcome, people stick to their own tempo regardless of their partner’s performance, like a drummer beating the song tempo and avoiding being dragged by the guitarist’s solo, or they synch with each other as best as possible, as two musicians playing an ostinato pattern. These two scenarios distinguish *emergent* and *planned* synchronization (Knoblich et al., 2011), where people synch involuntarily (‘entrainment’) or voluntarily with others, respectively. However, the interplay between the two processes and their reciprocal influence during visuomotor interactions is partially unknown.

The neurofunctional underpinnings of interpersonal synchronization (Repp, 2013)are still unclear as well. In a few fMRI studies (Fairhurst, Janata, & Keller, 2013; 2014), participants performed a finger-tapping task synching to a virtual partner programmed to adapt its frequency to the participants’ taps inducing different levels of coordination.

Results showed that the virtual partner’s adaptivity modulated the brain activity both in cortical (e.g., the dorsomedial prefrontal cortex and the superior frontal gyrus) and subcortical (e.g., the hippocampus and the insula) areas. However, the generalizability of these results is limited given that virtual partners are governed by pre-specified algorithms, which behavior does not fully reproduce the one typical of real-life visuomotor interactions between human agents. For instance, participants rush during dyadic tasks revealing worse performance than in solo conditions (Wolfe & Knoblich, 2022).

Another fMRI study (Silfwerbrand et al., 2022) explored leader-follower dynamics between pairs of participants during an audio-motor coordination task. The results revealed the prominent role of the cerebellum during both leading and following, suggesting it may be a key brain structure supporting interpersonal coordination. However, these results are confined to the audio-motor domain and cannot be extended to visuomotor interactions, in which spatiotemporal information processed through visual pathways plays a crucial role. Visuomotor interactions entail the non-trivial issue of discerning planned from emergent processes. Whether people synchronize voluntarily or involuntarily, sensory information about their partner’s movements is the same. Since observing a compatible movement elicits entrainment (Hove, Spivey, & Krumhanls, 2010; Oullier et al., 2008), a crucial question is whether voluntary co-regulation (planned synchronization) may override entrainment (emergent synchronization) in some circumstances and what neurophysiological processes mediate such an effect.

We recently tackled the above issue by using a finger-tapping task (Uccelli, Sacheli, & Paulesu, 2023), in which pairs of participants observed their partner’s hand while (i) synching with each other to replicate the tempo together, or (ii) resisting synchronization and focusing on individual performance. We introduced a tempo aftereffect within the dyad so that the tempo was the same but perceived slightly differently by participants who were unaware of this manipulation. This allowed us to estimate ‘tempo contagion’, that is, a measure of the timing deviation toward the partner’s tapping. We observed similar contagion effects when either participants synched or resisted, indicating that entrainment survived voluntary corrections. However, tempo contagion did not correlate within participants between the two conditions, suggesting two (potentially) different neurocognitive mechanisms underpinning emergent and planned synchronization. We adapted our previous paradigm to an fMRI setting to address this hypothesis.

### Aims

We investigated the neurofunctional underpinnings of emergent and planned synchronization by asking pairs of participants to synch with each other or to resist interpersonal synchronization during fMRI (see Methods). We expected to observe tempo contagion in both scenarios, associated with the activation of cortical and subcortical brain structures typical of synchronization tasks (including primary sensorimotor areas, the cerebellum, and higher-order visual areas involved in biological movement observation). However, we hypothesized that if emergent and planned synchronization are partially dissociated, the relation between behavior (as measured by the ‘tempo contagion’ effect) and brain activity should dissociate. Finally, we conducted a connectivity analysis to investigate whether task instructions may modulate the functional connectivity between the brain areas involved in the task.

## Methods

### Ethics

The Milano-Bicocca University Ethics Committee approved the experimental protocol [n°735]. Participants signed written informed consent. The study was conducted in accordance with standards of the Code of Ethical Principles for Medical Research Involving Human Subjects of the World Medical Association (Declaration of Helsinki), of the Italian Board of Psychologists (https://www.psy.it/codice-deontologico-degli-psicologi-italiani), as well as the Ethical Code for Psychological Research of the Italian Psychological Society (https://aipass.org/node/11560).

### Sample size determination

We conducted an a-priori power analysis to determine the adequate sample size to replicate the tempo contagion effect. In our previous work (Uccelli, Sacheli, & Paulesu, 2023), we observed an average tempo contagion of ∼2% (st. dev. ∼2.6). We calculated power assuming a similar average under three hypothetical scenarios where the group st. dev. (i.e., the inter-participant variability) was: i) equal, ii) 3.5 (+30%), or iii) 3.9 (+50%). As we expected a positive tempo contagion average (see below), we estimated power with a right-tailed *t*-test (i.e., not testing for a negative average). Power was calculated with the *pwr* package in R (Champley et al., 2018) with an increasing *n.* For *n* = 30, power estimates were .99, .92, and .86 for the three scenarios, respectively. Thus, we deemed *n* = 30 as adequate to replicate our tempo contagion estimates in the scanned participants’ sample (see next section).

### Participants

Sixty volunteers from the Milano-Bicocca community formed 30 dyads (42 females; mean age: 24.5 range: 18-33) not necessarily matched for gender (f-f: 13, f-m: 16, m-m: 1). They were all right-handed (by self-report), had normal hearing and normal or corrected to normal vision, had no history of neurological diseases, had no professional musical expertise, and were naïve as to the purpose of the experiment. Students recruited through a local SONA system (https://milano-bicocca.sona-systems.com) received formative credits. Thirty volunteers entered the fMRI scanner as experimental subjects (‘IN’; 16 females), whereas the other thirty stood beside the cot as partners (‘OUT’). Three IN participants were excluded for the following reasons: one was left-handed and therefore considered a pilot; one revealed an excessive number of movement artifacts; and one revealed a loss of signal in the right prefrontal area because of technical issues. Accordingly, those participants were excluded and the final sample involved 27 dyads.

### Stimuli

Stimuli were .mp3 files of a cowbell-like click (10 ms; 4800 Hz) generated in Audacity (https://www.audacityteam.org). For IN participants, three inter-onset-intervals (IOIs) were used as target tempi: 400, 500, and 600 ms (150, 120, and 100 bpm). For OUT participants, three IOIs centered around the target tempi of IN participants were used as distractor tempi: one -20 ms faster and one +20 ms slower (“incongruent” tempi), and one medium tempo equal to the target (“congruent” tempi, which served as baseline). We have chosen 20 ms as it is about double the 2% average tempo contagion observed in our previous work (see the power analysis above). These three triplets (**Fig.1a**) allowed us to estimate tempo contagion (see below). Participants were unaware that tempi could be either the same or different between partners. Each tempi sounded for 6 times during the synchronization phase (see *Procedure* below).

**Figure 1.**
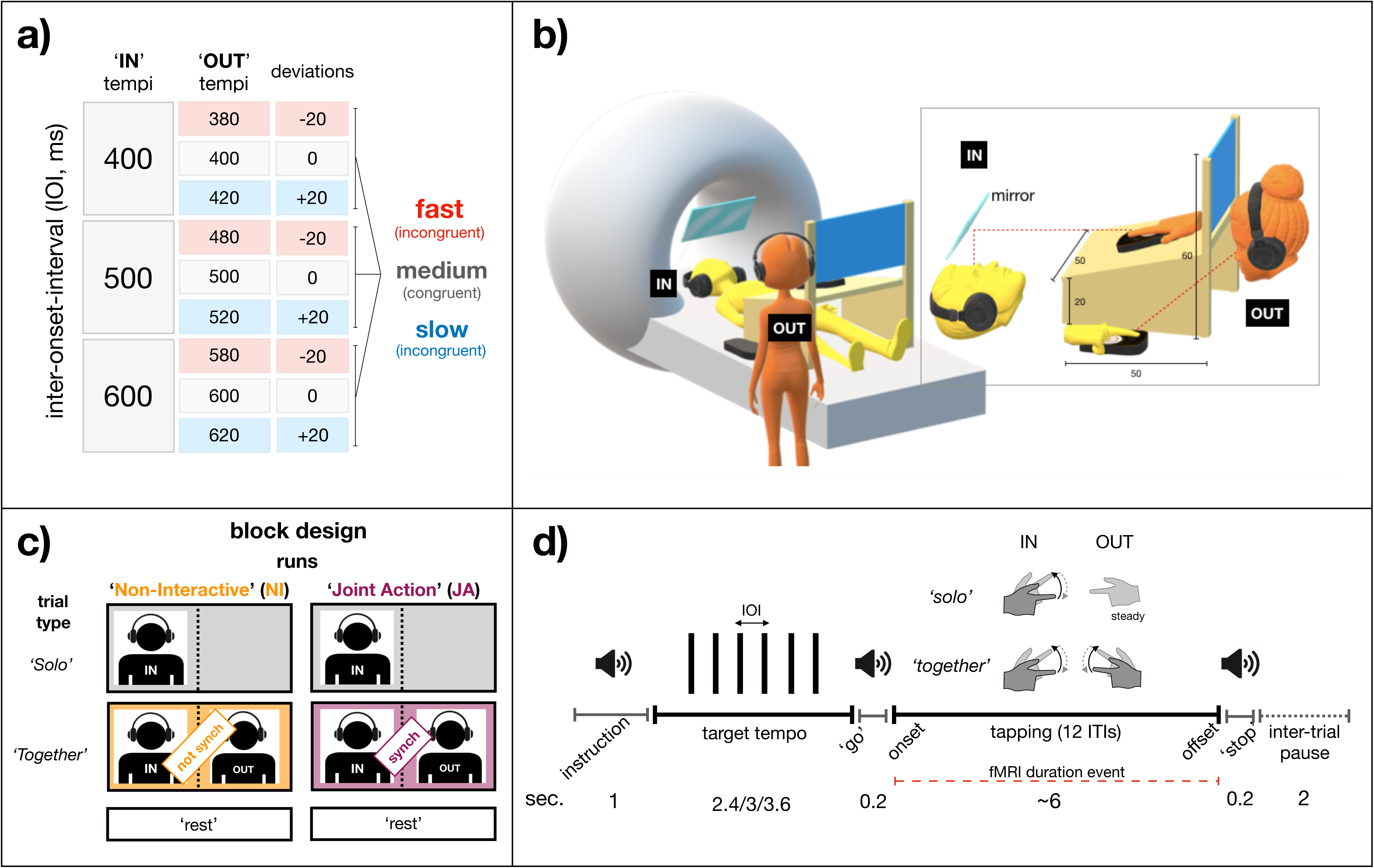
Methods overview. a) Stimuli. ‘IN’ fMRI participants listened to three target tempi (first column). For each, ‘OUT’ fMRI participants listened either to a slower (+20ms), a faster (-20ms), or an equal distractor tempo, yielding three triplets of tempi (middle column). This structure was reduced by re- expressing the data as deviations from the medium-medium pair of tempi (i.e., no effect baseline), isolating the ± 20 ms differences (third column). This re-expression yielded three triples centered around zero reducing inter-participant variability and improving estimation of tempo contagion. **b) Setup.** A mirror on the head cage allowed IN participants to observe the OUT participant’s hand leaning on a custom-made table above the cot. OUT participants stood beside the fMRI cot, observing the IN partner’s hand in front of them. Two response boxes collected tapping data. **c) Experimental design.** The two runs (‘Non-interactive’, NI, and ‘Joint Action’, JA) consisted of ‘Solo’ (only IN participants tapping), ‘Together’ (both participants tapping), and rest trials. Runs were identical except for Together trials in which the instruction was not to synch (NI) or synch (JA) with the partner’s tapping. **d) Trial timeline.** Participants listened to the instruction relative to the trial type, then the appropriate tempo sounded six beats. Once ceased, participants tapped maintaining the tempo according to the instruction until a stop sound signaled the trial’s end. See text for details.

### Setup and apparatus

**Fig.1b** depicts the experimental setup. OUT participants stood beside the fMRI cot, seeing the IN participants’ right hand. An fMRI-compatible table on the cot held the OUT participants’ hand. IN participants were able to see the OUT participants’ hand through a mirror placed on the head cage. A panel fixed on the back of the table prevented OUT participants from seeing their hand, as well as focusing the IN participant’s vision on the OUT participant’s hand. Two fMRI-compatible response boxes (Resonance Technology Inc., Northridge, CA, USA) were used to collect tapping data. Both participants wore sound-proofing headphones to listen to tempi and communicate with the control room.

### Experimental conditions and procedure

Participants performed an unpaced finger-tapping task while keeping their gaze on the partner’s hand. The general instruction was “synchronize to the target tempo and, once ceased, maintain it at best by tapping your right index until the trial ends”. Participants completed two runs: a ‘Non-interactive’ (NI) and a ’Joint Action’ (JA) run. Each run included the following three types of trials (**Fig.1c**):

1. ‘*Solo*’. IN participant listened to the target tempo and replicated it individually while observing the hand of the OUT partner, who was instructed not to take any action and to wait for Together trials. Solo trials for IN participants were included to subtract the neural activity of individual synchronization from that of interpersonal synchronization (see fMRI analysis below). Solo trials for OUT participants were unnecessary and therefore not included.
2. ‘*Together*’. Both participants listened to their appropriate tempo and replicated it according to the instruction: in the NI run, the instruction was to ‘replicate the tempo at best trying not to fall in synch with the partner’s tapping’. Conversely, in the JA run, the instruction was to ‘synchronize with the partner’s tapping to maintain jointly the tempo as best as possible’.
3. ‘*Rest’*. No tempo sounded. Participants relaxed waiting for the next trial.

Note that NI and JA Together trials provided identical visuomotor information and differed only in the type of synchronization instruction. NI and JA runs were counterbalanced across participants.

Dyads were pre-instructed by completing 10 practice trials with randomly chosen tempi for each condition on a PC prior to the fMRI experiment. In addition, other 5 practice trials were run in the fMRI room before initiating the experiment. **Fig.1d** depicts the trial timeline. At the beginning of each trial, an acoustic instruction informed the dyad of the condition (‘Solo’ or ‘Together’). Next, tempi sounded 6 times and dyads synchronized to them (‘synchronization’ phase), then the ‘go’ signal sounded. At this point, according to the condition, participants tapped (unpaced) maintaining the tempo until the ‘stop’ signal sounded signaling the trial’s end (‘tapping’ phase). The trial ended once the MATLAB script collected 13 taps (i.e., 12 intervals) from both participants; if the tapping phase exceeded a timeout of 12s, the trial ended and was considered an error. Rest trials lasted 12s to match the maximum trial time (i.e., 12s). Each run consisted of 63 randomized trials, including 18 Solo, 18 Together congruent (i.e., same tempo), 18 Together incongruent (9 fast and 9 slow distractors), and 9 rests . Each run lasted ∼13 min separated by a brief pause, plus ∼8 min for acquiring the structural (T1) image. Scanning lasted ∼40 min in total.

### Behavioural analysis

We analyzed the inter-onset-interval (ITI), the difference in time between two consecutive taps (tap_n_ - tap_n-1_), for both IN and OUT participants. From ITIs, we estimated tempo contagion effects (see below) to quantify the magnitude of temporal entrainment.

#### Data validation

We visually inspected the individual ITI raw distributions as a function of IOIs for detecting outliers. First, we considered ITIs < 150 and > 1000 ms as extreme values and therefore removed. Second, we applied individually a filter to detect ITIs ± 250 ms of the actual IOI in accordance with our previous work (Uccelli, Sacheli, & Paulesu, 2023). In accord with this criterion, 7.3% ITIs were excluded in Solo trials, while 7% and 8.3% ITIs for IN and OUT participants, respectively, were excluded in Together trials. Third, we checked normal assumptions in the ITI distributions to apply a transformation in case of marked asymmetry. The distribution was right-skewed and platykurtic for both participants (IN: skewness = 0.33, kurtosis = 3.9; OUT: skewness = 0.44, kurtosis = 3.86). A *boxcox* procedure returned a λ value of ∼0.5 (i.e., a sq. rt. transformation, see Osborne, 2010) for both distributions, indicating that the square root transformation could improve the distributions (see Osborne, 2010). The transformation made distributions more symmetrical (IN: -0.07; 4.1; OUT: 0.06; 3.8). For the sake of interpretation, we elevated to square individual means before the analysis.

#### Estimation of tempo contagion

We expected participants to drift ITIs toward each other in incongruent trials. For instance, IN participants should tap faster (or slower) when OUT participants tap faster (or slower), compared to trials in which the partner heard congruent tempi (baseline) (see **Fig.1a**). Inter-individual differences (e.g., the speeding up tendency in dyadic tasks called ’rushing’ effect, Wolfe & Knoblich, 2022; Wolf et al., 2019) may mask subtle temporal adjustments, making difficult estimating tempo contagion. To bypass this issue, we performed a centering variable procedure (Enders & Tofighi, 2007) to reduce inter-individual differences. For each triplet of tempi (for instance, 480, 500, and 520) we re-expressed data as deviations by subtracting congruent averages from each ITI datapoint, obtaining positive and negative values relative to zero, signaling slower or faster tap, respectively. After this re-expression, the three triplets of means consisted of a faster (-20 ms) and a slower (+20 ms) means around a common zero (see third column in **Fig.1a**). This re-expression isolated tempo contagion, yielding comparable estimates across participants, conditions, and targets. For the sake of clarity, given a 500 ms tempo for the IN participant and a 480 ms tempo for the OUT participant, we expected the former to show a negative (faster) deviation and the latter a positive (slower) deviation, compared to when they listened both to the 500 ms baseline.

Last, we estimated a unique tempo contagion effect as follows: i) computed the averages of slow and fast ITI deviations, ii) computed the difference between such averages, and then iii) expressed the difference as a percentage to the appropriate pair of medium tempi (baseline), as summarized by the following formula in accordance to our previous work (Uccelli, Sacheli, & Paulesu, 2023; see also Uccelli & Bruno, 2024):

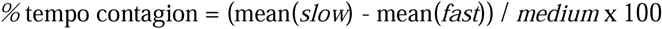

where the numerator is the difference between slow (+20 ms) and fast (-20 ms) average means, and the denominator is the appropriate medium (baseline) mean. Positive % values signal tempo contagion effects. The formula was applied individually in both NI and JA runs.

#### Analysis of tempo contagion

Percent tempo contagion effects entered into a 2x2 mixed ANOVA, with ‘Condition’ (NI/JA) as the within-subjects factor term and ‘Participants’ (IN/OUT) as the between-subjects factor term. If the two factors did not modulate tempo contagion, the model should return no significant main or interaction effects. Testing was run using the *ezANOVA* function of the R *ez* package (Lawrence & Lawrence, 2016).

### MRI data acquisition and analyses

#### Data acquisition

Whole-brain functional images were acquired using a 3.0-T Ingenia CX scanner (Philips). Gradient-echo T2*-weighted transverse echo-planar images (EPI) were acquired (scan parameters: TR = 2000 ms, TE = 30 ms, 35 transversal slices, descending not interleaved acquisition, 4 mm slice thickness, with no inter-slice gap, FA = 75°, FOV = 240 mm, matrix size = 80 × 80). A high-resolution T1-weighted anatomical scan (TR = 8.1 ms, TE = 3.7 ms, 207 sagittal slices, FOV = 256 mm, matrix size = 256 × 256, FA = 8°, Inversion Time (TI) = 1000 ms) was also acquired. Each run was planned to acquire a maximum of 460 scans. The expected number of scans per run was ≈ 440, but we extended to a maximum of 460 to ensure that the run did not end before participants completed all trials. If dyads completed the task beforehand, the run was stopped, and the last scan was discarded before the preprocessing steps to avoid potential distortions due to the forced interruption. On average, 437 scans (st. dev. = 21.7) were acquired per run.

#### Task functional activation analysis

Images were converted from DICOM to NifTI format in mriconvert (https://neuro.debian.net/pkgs/mriconvert.html). Successive analyses were performed in MATLAB (R2019b, MathWorks) using Statistical Parametric Mapping (SPM12, Wellcome Department of Imaging Neuroscience).

#### Preprocessing

First, scans were realigned and unwrapped to account for any movement during the experiment and to reduce magnetic field distortion effects. Second, unwrapped images were coregistered with a T1-weighted structural image of each participant, which was then segmented and stereotactically normalized into the SPM12 template (tmp.nii) to allow for group analyses. Third, deformation fields used for T1 segmentation were applied to coregistered functional scans; here, the data matrix was interpolated to produce voxels 2x2x2 mm in dimension. Last, the normalized images were smoothed using a Gaussian filter of 10 × 10 × 10 mm to improve the signal-to-noise ratio and allow for valid application of the family-wise error rate (FWER) correction at the cluster-level in the second-level (i.e., group-level) univariate analyses, as recommended by (Flandin & Friston, 2019). We used Artifact Detection Tools (ART, Whitfield-Gabrieli, https://www.nitrc.org/projects/artifact_detect) to identify outlier scans for each participant. Time- points were marked as outliers when scan-to-scan variations in the global signal exceeded 3 st. dev. from the mean and when the compounded measure of movement parameters exceeded 1 mm scan-to-scan movement. On average, excluded volumes were 2.14%. Only one participant who showed 135 outliers (31.1%) was excluded from the sample.

#### Statistical analyses

We performed a two-step statistical analysis based on the general linear model. The blood oxygen level-dependent (BOLD) signal associated with each experimental condition was analyzed by convolution with a canonical hemo-dynamic response function (Worsley & Friston, 1995).

Global normalization was applied. The time series was high-pass filtered at 128s and prewhitened by means of an autocorrelation model AR(1).

In a block design, the event onset corresponded to the tapping onset, and the event duration corresponded to the difference between the go signal and the moment at which 13 taps were recorded from both participants (see the dashed red line in **Fig.1d**). Separately for the JA and NI runs, these events entered the analyses to estimate the BOLD signal associated to the following conditions of interests: Solo and Together. Rest trials were not modeled, following (Pernet, 2014). Regressors of non-interest modeled experimental confounds, including the signal associated with: i) the onset of targets, ii) the onset of the instruction, go, and stop signals, iii) the realigning parameters calculated in the prepossessing step, and iv) errors (that is, trials either exceeding the timeout or having an anomalous duration compared to other trials, < 1%).

At the first-level (individual) analysis, we computed fixed condition-specific effects for each participant. We modeled the simple effect of each condition of interest (Solo and Together, separately for the NI and JA runs). Such contrasts allowed us to determine both individual- and dyadic-specific activations and their difference.

At the second-level (group) analysis, we computed a 2x2 full-factorial design including Task (Solo vs. Together) and Condition (NI vs. JA) as within-subject factors. As a preliminary analysis, we checked for the presence of any significant difference between the Solos in the JA and NI runs (to ensure they did not differ) and explored the simple effect of Solo that served as a reference point. Then, we explored the main effect of Condition (Solo > Together, Together > Solo, see Supplementary Materials) and the interaction effect (NI(Together > Solo), JA(Together > Solo)). We also planned a conjunction analysis to describe the brain areas co-activated during NI and JA runs. All contrasts were masked with the main effect of the task (contrast: 1 1 1 1, p < .05 uncorr.) to only include significant task-related activations.

The reported results apply a family-wise error rate (FWER) correction for multiple comparisons at the cluster-level (p < .05). The cluster-wise correction was applied to data having applied a 10x10x10 Gaussian smoothing and at p < .001_uncorr_ at the voxel-level, as recommended by Flandin & Friston (2019).

#### Linear regression between % tempo contagion and BOLD signal

We planned to extract the effect of interest (JA or NI task-related activation > Solo) from the active clusters during NI and JA Together trials to test their association with the % tempo contagion effect. The average *y*-values for each cluster were extracted by using the *MarsBaR* toolbox (Brett et al., 2022). We added the values extracted from each cluster as continuous predictors in a series of linear models fitting % tempo contagion effects and having ‘Condition’ (NI/JA) as an additional factor. We expected task-specific activations extracted from different clusters to modulate tempo contagion effects differently in the JA and NI trials, as possibly shown by a significant interaction effect. To evaluate such potential interactions, we computed 95% CIs around the slopes judging whether they covered zero.

#### Task functional connectivity analysis

Functional connectivity analyses were performed in MATLAB using the software CONN 22.a (Whitfield-Gabrieli & Nieto-Castanon, 2012) and SPM12.

#### Preprocessing and denoising

Scans were preprocessed according to the following steps: realignment, slice timing correction, outlier detection (ART, see above), indirect segmentation and MNI-space normalization, and smoothing (10x10x10 mm). Scans were denoised using a standard pipeline including the following regressors of potential confounding effects: white matter and CSF timeseries, motion parameters and their first-order derivatives, task activation effects, outlier scans, and linear trends within each run. A high-pass 0.008 Hz frequency filter was applied to the BOLD time series. No low-pass filter was applied. For additional information, see the Supplement materials online.

#### First-level analysis

Seed-based functional connectivity maps were estimated characterizing the spatial pattern of the whole-brain functional connectivity with the seed areas, as defined by the brain clusters that correlated with % tempo contagion estimates in the task activation analyses. Functional connectivity strength was represented by Fisher-transformed bivariate correlation coefficients estimated separately for each seed area and target voxel, modeling the association between their BOLD signal timeseries.

#### Second-level analysis

Linear regressions were performed to test whether the % tempo contagion was associated with the whole-brain functional connectivity of the selected seeds. In one regression model, we used the NI % tempo contagion to predict the whole-brain functional connectivity strength of the seed areas during the NI condition. In a separate regression model, the JA % tempo contagion was used to predict the functional connectivity strength of the seed areas during the JA condition.

Voxel-level hypotheses were evaluated using multivariate parametric statistics with random effects across subjects and sample covariance estimation across multiple measurements. Results were thresholded at p < .001_uncorr_ at the voxel-level and p < .05_FWE_ at the cluster-level.

## Results

### Behavioral results

#### Solo trials

We checked whether ITIs of IN participants did not differ between NI and JA runs. Group averages with standard error of the mean (SEM) were almost equal across the three IOIs [NI vs. JA: 392 ± 3 and 391 ± 4 ms; 470 ± 4 and 470 ± 4 ms; 550 ± 5 and 545 ± 5 ms].

Although ITIs scaled along the IOIs, on average, participants anticipated the actual tempo, in accord with rushing effects. Visual inspection confirmed that the group averages did not differ between conditions (see Fig.S2 in the supplemental materials and the relative testing).

#### Together trials

**Fig.2a** presents individual and group ITI means as a function of tempi in both runs (left: NI; right: JA). Each point can be compared to zero vertically or horizontally for IN and OUT participants, respectively. Dyads underestimated the 500 and 600 ms tempi (see distances between SEMs from orthogonal crosses), consistent with rushing effects. Given that rushing may potentially mask fine co-adjustments in time, we computed deviations from the baseline average to isolate tempo contagion (see Methods for details).

**Figure 2.**
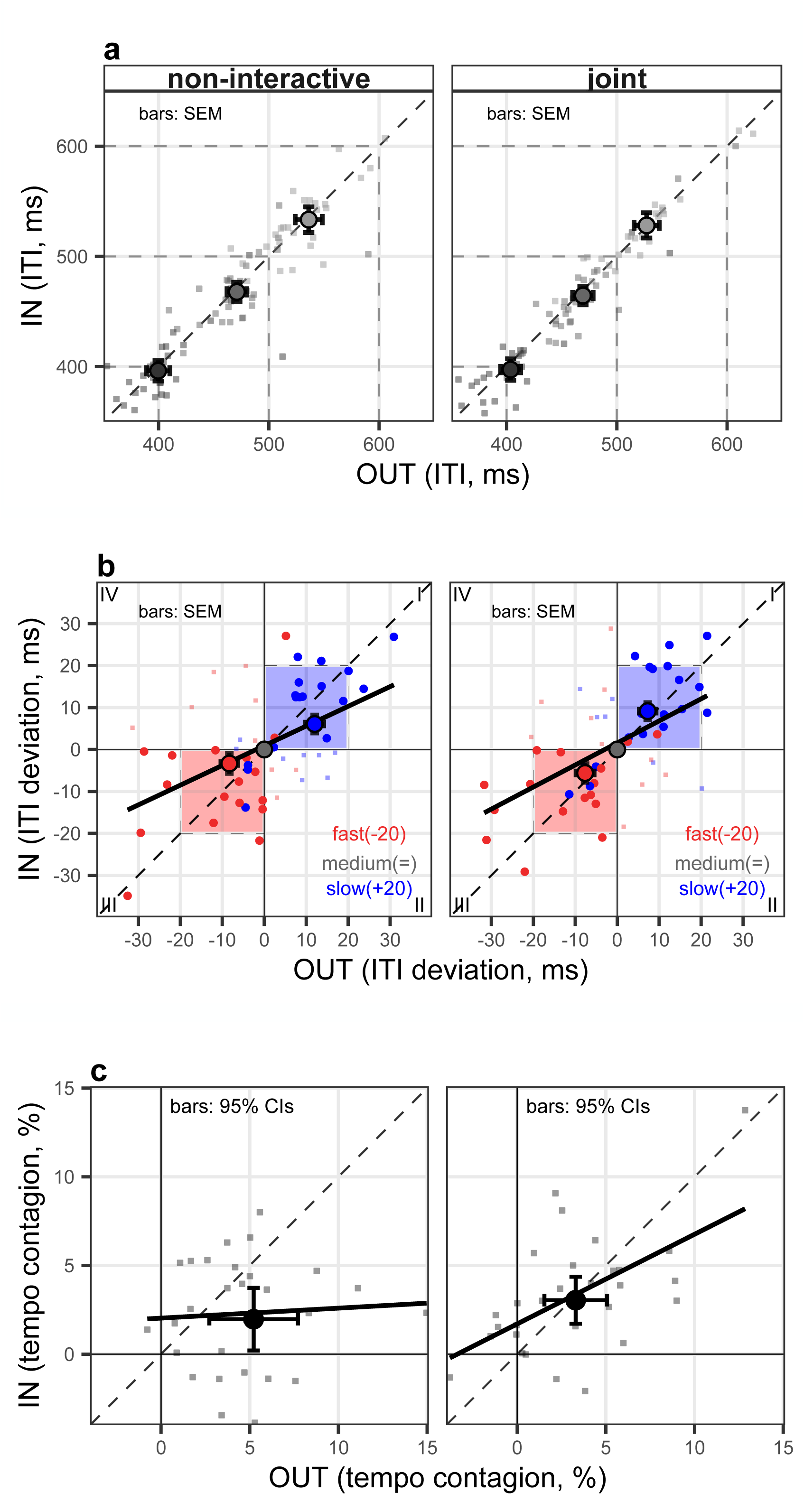
Behavioral results. Individual (small points) and group (large points) data of ‘IN’ fMRI participants (y-axes) are plotted against ‘OUT’ participants (x-axes) in the two Together ‘Non-interactive’ and ‘Joint Action’ conditions (columns). **a) Inter-tap-interval (ITI).** ITIs (in absolute values) scaled on the actual tempi, but dyads rushed at 500 and 600 ms (group averages do not cover the orthogonal crosses). **b) Estimation of tempo contagion.** ITIs re-expressed as deviations from the medium-medium mean (baseline, grey points). Group averages on the y-axis indicate that IN participants tapped faster or slower when the OUT partner tapped for a faster (red points, -20 ms) or slower (blue points, +20 ms) distractor tempo. Conversely, group averages on the x-axis do not lie on the black segments (i.e., the actual distractor tempo) but are closer to the baseline average, indicating that OUT participants drifted tapping as well. Colored areas: the ranges in which ITIs are expected if both participants drift toward each other. Thick black line: linear regression obtained by averaging parameters from individual fits. **c) Percent tempo contagion.** Both IN and OUT participants showed similar % tempo contagion effects across conditions (group averages’ CIs do not cross zero lines). See Methods for the formula description. Thick black lines: linear regression fitting % tempo contagion effects of IN and OUT participants.

**Fig.2b** presents the individual and group ITI deviation means. For instance, for a target and a distractor of 500 and 520 ms, respectively, participants should drift tapping toward the middle (∼510). Indeed, most of the fast tempi (red) led to faster ITIs (quadrant III) and most of the slow tempi (blue) to slower ITIs (quadrant I): group averages fall within the two colored regions in which tapping was expected to balance between partners.

**Fig.2c** presents individual and group % tempo contagion effects (see Methods for the formula description). Group averages were similar across participants and conditions. The mixed ANOVA (see Methods for details) revealed no main effect of Role (IN/OUT) [F(1, 52) = 3.73, p = .06, η^2^ = .036] and Condition (NI/JA) [F(1, 52) = 0.001, p = .98, η^2^ = .06], nor the Role x Condition interaction effect [F(1, 52) = 2.66, p = .11, η^2^ = .02], but a significant effect of the intercept [F(1, 52) = 8.3, p < .001, η^2^ = .45]. Thus, % effects were different from zero (see CIs in **Fig.2c** not crossing zero lines) but not modulated by experimental manipulations. Moreover, % effects correlated between IN and OUT participants in the JA run [*r*(25) = .59, CI(.27, .79); p = .001] but not in the NI one [*r*(25) = .21, CI(-.18, .54); p = .29]. Last, % effects did not correlate between NI and JA conditions either for IN [*r*(25) = -.0003, CI(-.37, .37); p = .29] or OUT [*r*(25) = .08, CI(-.31, .45); p = .29] participants.

### fMRI results

#### Preliminary analysis on the Solo network

We checked whether Solo trials did not show different activations between NI and JA runs. The two ‘Solo NI > Solo JA’ and ‘Solo JA > Solo NI’ linear contrasts revealed no difference. Next, we calculated the simple effect of ‘Solo’ to investigate brain regions specifically active during individual tapping. Such brain regions included (see **Fig.3a**, grey clusters) the bilateral precentral gyrus, left postcentral gyrus, left supplemental motor area, left insula, left pallidum and putamen, and lobules VI and VIII of the right cerebellum (for details, see Table S1 in the Supplemental).

**Figure 3.**
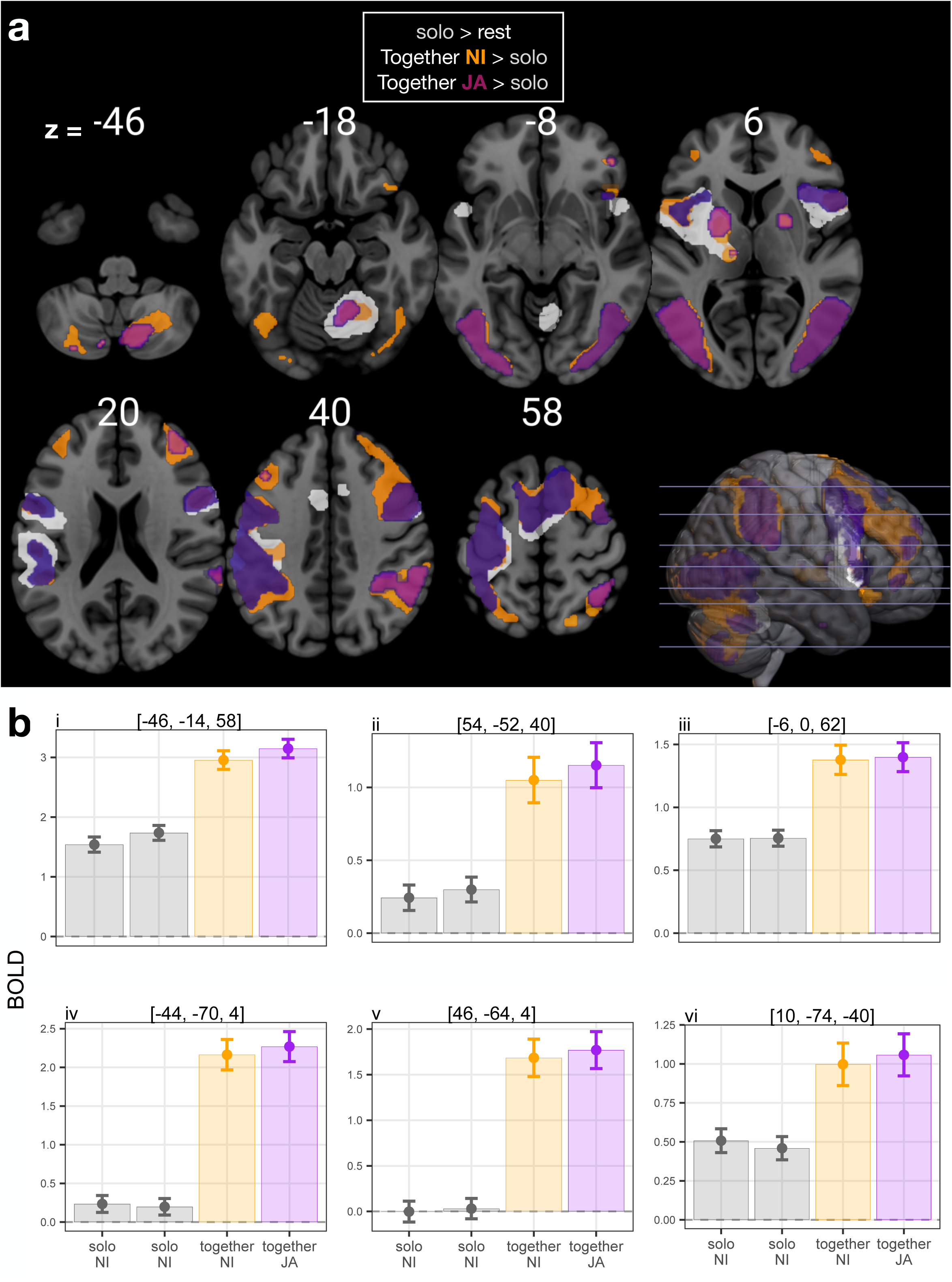
Results of the activation task. **a)** results from the Task (Solo vs. Together) x Condition (Non- Interactive, ‘NI’, vs. Joint Action, ‘JA’) ANOVA. Grey clusters represent the simple effect of Solo in each run (NI vs. JA). Brain regions active during Together trials more than Solo trials are depicted separately for the NI (orange) and JA (purple) runs. Data is plotted by applying a p_uncorr._ < .001 threshold at the voxel level and a p_FWER-corr._ < .05 threshold at the cluster level (see Table S2 in the supplemental materials for details on MNI coordinates). **b)** Average BOLD signal of the ‘Together > Solo’ linear contrast at the local maxima of: left (i) and right (ii) precentral gyrus (PrG), left supplemental motor area (SMA, iii), left (iv) and right (v) lateral occipitotemporal cortex (LOTc), and right cerebellum (vi). Barplots show beta values with 95% CIs. Note that the overlapping of orange and purple CIs signal no interaction effects (i.e., NI(Together > Solo) and JA(Together > Solo) linear contrasts).

*Together network -“Together > Solo”*

**Fig.3a** presents the active brain clusters of the ‘Together > Solo’ linear contrast, split in orange and purple for the NI and JA runs, respectively. Overall, the BOLD signal increased in the same clusters observed in Solo (see orange and purple bars compared to grey ones in **Fig.3b**, which plots the simple effects of each task compared to the implicit baseline). Both the NI and JA Together tasks activated two large bilateral clusters in the lateral occipitotemporal cortex (LOTc), which were silent during Solos, and a smaller portion of the right cerebellum was more active in both NI and JA conditions. However, the ANOVA revealed no significant difference between the ‘NI (Together > Solo)’ and ‘JA (Together > Solo)’ linear contrasts (i.e., the interactions; see the orange and purple bars in **Fig.3b**; see also Tab. S2 in the Supplemental for detailed information). Last, the ‘Solo > Together’ linear contrast did not reveal significant activations.

*Conjunction analysis - “NI(Together > Solo)* ∩ *JA(Together > Solo)”*

Given extensive overlapping activations in the social conditions, we proceeded with a conjunction (inclusive) analysis [NI(Together > Solo) ∩ JA(Together > Solo)] to restrict the regression analysis with % tempo contagion (see next paragraph) to the brain areas co- active in both tasks. The following six clusters were significantly activated in both conditions (**Fig.4a**): the right cerebellum (lobule VI), the bilateral LOTc, the right frontal and parental clusters, and a large left frontoparietal cluster. **Table 1** reports the MNI coordinates of such clusters and the relative significance (p < .001_uncorr_ at the voxel-level and p < .05 FWER_corr._ at the cluster level).

**Figure 4.**
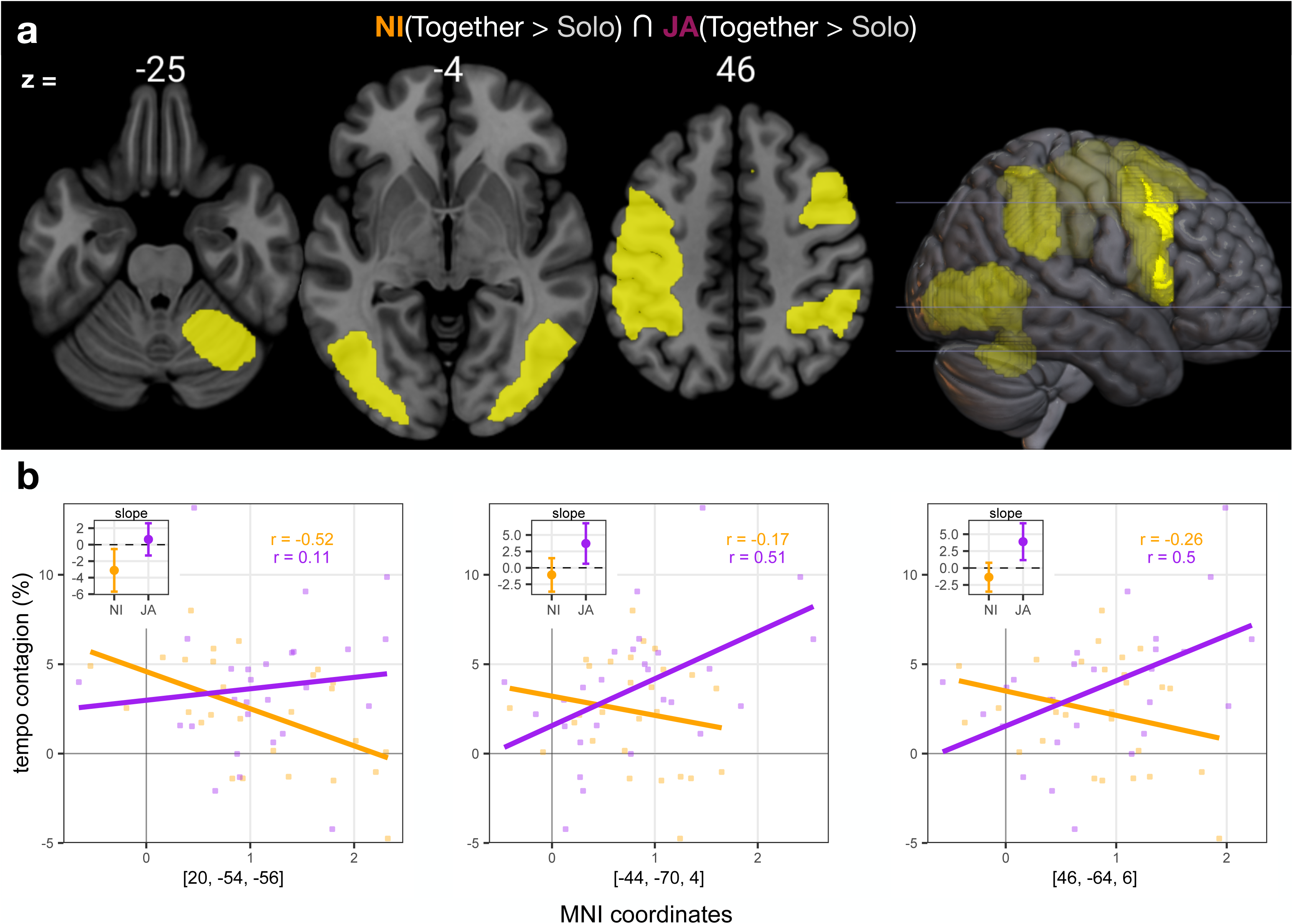
Results of the Conjunction analysis. **a)** clusters from the NI (Together > Solo) ∩ JA (Together > Solo) conjunction. Data is plotted by applying a p_uncorr._ < .001 threshold at the voxel level and a p_FWER-corr._ < .05 threshold at the cluster level. **b)** Linear regression between the individual % tempo contagion (y-axis) and the average BOLD signal (x-axis) of the cluster computed with the *MarsBar* toolbox (left plot: right cerebellum; middle and right plots: the bilateral LOTc). Each point represents a different IN participant. Inset plots: 95% confidence intervals (CIs) around slopes extracted from the linear model (see text for details). CIs not covering zero (dashed line) indicate that BOLD signal modulated % effects. Orange: Non-interactive (NI); purple: Joint Action (JA). Note that the regressions with the right parietal cluster, and right and left frontal clusters are not shown since they did not reveal differences in slope, but plots are presented in Fig. S2 in the supplemental online.

**Table 1.** Conjunction analysis “NI (Together > Solo) ∩ **JA (Together > Solo)”.** X, Y, and Z are the stereotactic coordinates of the activations in the MNI space. All reported voxels (p < .001_uncorr_) are included in clusters surviving the FWER correction at the cluster-level. A maximum of 16 coordinates (local maxima) per cluster have been reported, each placed at 4 mm apart, as reports in SPM12. (*) Z-scores statistically significant also after whole brain FWER correction (p < .05) at the voxel-level (FWER: family-wise error rate; MNI: Montreal Neurological Institute).

| Brain area<br>(Brodmann) | Left Hemisphere |  |  |  | Right Hemisphere |  |  |  |
| --- | --- | --- | --- | --- | --- | --- | --- | --- |
|  | X | Y | Z | Z-score | X | Y | Z | Z-score |
| **Left occipital cluster**<br>k = 1408, $p_{\text{FWE-corr.}} = .001$ | | | | | | | | |
| Medial occipital gyrus (37) | -44 | -70 | 4 | >8* | -- | -- | -- | -- |
| Inferior occipital gyrus (18) | -28 | -94 | -8 | 6.77* | -- | -- | -- | -- |
| **Right tempo-occipital cluster**<br>k = 1500, $p_{\text{FWE-corr.}} = .001$ | | | | | | | | |
| Medial occipital gyrus (37) | -- | -- | -- | -- | 46 | -64 | 6 | >8* |
| Medial occipital gyrus (19) | -- | -- | -- | -- | 38 | -88 | 2 | 6.74* |
| **Right parietal cluster**<br>k = 1187, $p_{\text{FWE-corr.}} = .002$ | | | | | | | | |
| Inferior parietal gyrus (40) | -- | -- | -- | -- | 56 | -52 | 40 | 5.12* |
|  | -- | -- | -- | -- | 54 | -52 | 44 | 5.03* |
|  | -- | -- | -- | -- | 42 | -48 | 52 | 4.58* |
|  | -- | -- | -- | -- | 46 | -46 | 52 | 4.57* |
|  | -- | -- | -- | -- | 32 | -50 | 48 | 4.56* |
| Angular gyrus (40) | -- | -- | -- | -- | 62 | -52 | 28 | 4.41* |
| Supramarginal gyrus | -- | -- | -- | -- | 60 | -38 | 28 | 4.13 |
|  | -- | -- | -- | -- | 66 | -40 | 26 | 4.04 |
| **Fronto-parietal cluster** |  |  |  |  |  |  |  |  |
| k = 8423, p<sub>FWE-corr.</sub> < .001 |  |  |  |  |  |  |  |  |
| Post-central gyrus | -46 | -14 | 58 | >8* | -- | -- | -- | -- |
|  | -38 | -20 | 54 | >8* | -- | -- | -- | -- |
| Pre-central gyrus (6) | -46 | 0 | 52 | 6.94* | -- | -- | -- | -- |
| Inferior parietal gyrus (40) | -52 | -48 | 50 | 6.42* | -- | -- | -- | -- |
|  | -34 | -48 | 50 | 5.86* | -- | -- | -- | -- |
| Supplemental motor area (6) | -6 | 0 | 62 | 6.24* | 10 | 6 | 62 | 5.67* |
|  | -- | -- | -- | -- | 12 | 18 | 56 | 5.49* |
| Supramarginal gyrus (48) | -48 | -44 | 30 | 5.33* | -- | -- | -- | -- |
|  | -52 | -24 | 24 | 4.62* | -- | -- | -- | -- |
| Superior frontal gyrus (8) | -14 | 16 | 58 | 3.49 | -- | -- | -- | -- |
| Superior frontal gyrus (6) | -20 | 10 | 66 | 3.42 | -- | -- | -- | -- |
|  | -24 | 8 | 68 | 3.33 | -- | -- | -- | -- |
| Inferior frontal gyrus, pars opercularis (48) | -48 | 10 | 4 | 4.56* | -- | -- | -- | -- |
| Inferior frontal gyrus, pars opercularis (6) | -58 | 8 | 18 | 4.19 | -- | -- | -- | -- |
|  | -54 | 8 | 12 | 4.12 | -- | -- | -- | -- |
| **Right frontal cluster**<br>k = 1843, p<sub>FWE-corr.</sub> < .001 |  |  |  |  |  |  |  |  |
| Pre-central gyrus (6) | -- | -- | -- | -- | 48 | 2 | 50 | 6.04* |
| Inferior frontal gyrus, pars opercularis (6) | -- | -- | -- | -- | 56 | 12 | 12 | 4.48* |
| Inferior frontal gyrus, pars opercularis (44) | -- | -- | -- | -- | 52 | 10 | 20 | 4.14 |
|  | -- | -- | -- | -- | 52 | 10 | 26 | 4.11 |
| **Right cerebellum cluster**<br>k = 874, p<sub>FWE-corr.</sub> = .009 |  |  |  |  |  |  |  |  |
| cerebellum, VI | -- | -- | -- | -- | 20 | -54 | -24 | 5.88* |
| cerebellum, VI | -- | -- | -- | -- | 26 | -58 | -26 | 5.80* |

### Linear regressions between conjunction clusters and % tempo contagion

We tested the relation between % tempo contagion effects and the effect of interest (JA/NI > Solo) extracted from each cluster of the conjunction analysis (**Fig.4a)**. Since clusters were rather extended, the local maxima might not be representative of the cluster activity. Thus, we extracted the average BOLD *y*-values by using the *MarsBaR* toolbox. We computed a series of linear models fitting the % tempo contagion effects as a function of the condition and adding the values extracted with *MarsBaR* as a continuous predictor. Since % effects did not differ between conditions (see again **Fig.2c**), we were interested in evaluating interaction terms as they would indicate that the neurofunctional activations modulated % effects. For the cerebellum cluster, we observed a negative slope in the NI condition [t(50) = -2.7, p = .03] and a flat slope in the JA condition [t(50) = 0.6, p = .5]. For the right LOTc, we observed a positive slope in the JA condition [t(50) = 3.9, p = .005] but a flat slope in the NI condition [t(50) = -1.36, p = .2]. For the left LOTc, we observed again a positive slope in the JA condition [t(50) = 3.7, p = .02] but a flat slope in the NI condition [t(50) = -1.07, p = .4]. **Fig.4b** presents these regressions and the relative CIs around the slopes (inset plots). For the remaining clusters (i.e., the right parietal cluster, and right and left frontal clusters, not plotted in Fig.4b), the BOLD signal did not modulate tempo contagion (see Fig.S2 in the Supplemental materials).

### Task functional connectivity analysis

We sought to determine whether % tempo contagion effects modulated functional connectivity between conjunction clusters and the rest of the brain. Since contagion effects revealed a negative correlation in the right cerebellum in the NI condition and a positive correlation in the bilateral LOTc in the JA condition, we selected these clusters as seeds to restrict the analysis to the behavioral results observed on tempo contagion.

During the NI condition, the functional connectivity between the right cerebellum and a portion of the right LOTc (medial occipital gyrus) was negatively associated with NI % tempo contagion. Similarly, the functional connectivity between the right cerebellum and the right postcentral gyrus was negatively associated with NI % tempo contagion (**Fig.5a**; MNI coordinates in **Table 2**). Thus, the higher the NI % tempo contagion, the lower (more negative) the connectivity between the cerebellum and these two clusters (**Fig.5b**). No significant association emerged between NI % tempo contagion and the functional connectivity patterns of the right or left LOTc. During the JA condition, however, no significant association was observed between JA % tempo contagion and the functional connectivity patterns of the right cerebellum, or the right and left LOTc.

**Table 2.** Results of the functional connectivity with the percent tempo contagion in the Non-Interactive condition. Coordinates in MNI space, and Z-score of the resulting clusters are reported. (*) Z-scores statistically significant also after whole brain FWER correction (p < .05) at the voxel-level.

| Brain area<br>(Brodmann) | Left Hemisphere |  |  |  | Right Hemisphere |  |  |  |
| --- | --- | --- | --- | --- | --- | --- | --- | --- |
|  | X | Y | Z | Z-score | X | Y | Z | Z-score |
| **Right occipital cluster**<br>k = 904, $p_{FWE-corr.} < .001$ | | | | | | | | |
| Medial occipital gyrus (19) | -- | -- | -- | -- | 24 | -78 | 18 | -5.58 |
|  | -- | -- | -- | -- | 38 | -72 | 6 | -4.24 |
|  | -- | -- | -- | -- | 48 | -78 | 4 | -3.54 |
| Angular gyrus (39) |  |  |  |  | 44 | -76 | 4 | -4.18 |
| Superior occipital gyrus (7) | -- | -- | -- | -- | 24 | -76 | 38 | -3.53 |
| **Right frontal cluster**<br>k = 676, $p_{FWE-corr.} = .001$ | | | | | | | | |
| Pre-central gyrus (6) | -- | -- | -- | -- | 32 | -18 | 68 | -4.81* |
|  | -- | -- | -- | -- | 42 | -16 | 56 | -3.23 |
| Post-central gyrus (4) | -- | -- | -- | -- | 32 | -28 | 48 | -4.13 |

## Discussion

Real-life social interactions rely on exchanging visuomotor information, which is fundamental for achieving an individual or a shared synchronization. To the best of our knowledge, this work is the first to document the neurofunctional correlates of emergent and planned synchronization during visuomotor coordination. The results reveal a common neural network for both synchronization types, including the lateral occipitotemporal cortex (LOTc) and precentral gyrus, the right cerebellum, and the left postcentral gyrus. However, the behavioral temporal entrainment depended on the recruitment of different neural resources, indicating it depends on how visual and cerebellar brain regions process visuomotor information.

At the behavioral level, participants deviated their tapping toward the partner’s timing (**Fig.2b**), showing a similar tempo contagion magnitude between NI and JA conditions (**Fig.2c**). Thus, entrainment occurred regardless of the sought outcome, suggesting that emergent and planned processes might share similar neural mechanisms, in line with the behavioral results from our previous work (Uccelli, Sacheli, & Paulesu, 2023). However, % tempo contagion effects correlated between participants in the JA but not in the NI condition (**Fig.2c**); in contrast, this finding suggests that tempo contagion might stem from different neural mechanisms. During the NI task, participants were entrained to the partner’s timing to different degrees depending on the individual ability to resist entrainment. During the JA task, instead, participants ceded a similar amount of timing with each other given co-regulation strategies, leading to a similar tempo contagion. Depending on the social context (JA or NI) and the ensuing synchronization instructions, visuomotor information is exploited in different ways. In the NI task, the partner’s movement represents a source of interference, whereas in the JA task is fundamental to match as best as possible the partner’s timing. Therefore, different brain structures process visuomotor information in different manners, as it turned out in our data (see below).

At the neurofunctional level, Solo trials (**Fig.3a**, grey clusters) activated brain areas consistent with an ALE meta-analysis on finger-tapping tasks (Witt, Laird, & Meyerand, 2008). Among these areas, lobules VI and VIII of the right cerebellum are known to be involved in sequential finger movements (Stefanescu et al., 2013), broadly corresponding to the right hand somatotopy according to ALE meta-analyses (Boillat, Bazin, & van der Zwaag, 2020; var der Zwaag et al., 2013). Together (JA and NI) trials activated the same network during Solos, also in addition to two large (bilateral) visual clusters identifiable as the lateral occipitotemporal cortex (LOTc). LOTc is active during the observation of moving stimuli, including others’ actions (Zhang et al., 2022; Lingnau & Downing, 2015), a source of information lacking during Solo trials (as the OUT participant’s hand was still).

Yet, LOTc also includes a region (the extrastriate body area, EBA) selective to body parts, including the hand, partially overlapping with the MT/V5 cluster (Ferri et al., 2013).

Moreover, we observed a smaller portion of the right cerebellum (lobule VI; see orange and purple clusters compared to grey ones in **Fig.3b**) active in both NI and JA tasks. These findings indicate that the brain areas underlying interpersonal synchronization are approximately the same as individual synchronization, except for the recruitment of the LOTc. However, the NI and JA tasks did not show a voxel-by-voxel significant difference in the BOLD signal in any of the active clusters (see orange and purple bars in **Fig3.b**), suggesting that emergent and planned processes rely on the same neural network. It is not surprising not to observe differences in the activation task as entrainment survived during planned synchronization, as our tempo contagion analysis revealed.

The distinction between emergent and planned synchronization is partially unexplored. At least two meta-analyses on fMRI data (Teghil et al., 2019; Chauvignè et al., 2014) investigated external- and self-paced tapping, which, in a broader sense, resemble emergent and planned processes. In brief, externally-pace timing recruited more subcortical structures (e.g., the vermis) whereas internally-paced tapping more cortical structures (e.g., supplemental motor area). We did observe cortical and subcortical (e.g., the cerebellum) activations, but these activations were not specific activations for either the NI or JA task. Although these meta-analyses suggested a partial distinction between external- and self-paced tapping, it would be unwary to extend the same to a visuomotor context for three reasons. First, these meta-analyses focused on audio-motor tasks which are qualitatively different from visuomotor ones. Second, we demanded participants to voluntarily synch or not, each consisting of an active motor coordination strategy (different therefore from external-paced stimulation). Third, tempo contagion effects were similar in NI and JA tasks, indicating that entrainment occurred independently from the instruction.

We conducted a conjunction analysis (**Fig.4a**) to restrict the regression analyses with tempo contagion to the co-active clusters. As mentioned above, visuomotor information should promote or prevent tempo contagion depending on the instruction. The analysis supported this expectation on two grounds. First, the higher the BOLD signal in the LOTc, the higher the % tempo contagion effect, but only in the JA task (see **Fig.4b**). Second, the higher the BOLD signal in the cerebellum, the lower the % tempo contagion effect, but only in the NI (see **Fig.4b**). These findings suggest a two-fold implication. First, planned synchronization is driven by tracking the visuomotor information of the partner’s movement, probably due to motor mimicry (e.g., the timing of the tap peak amplitude; see

Kroger et al., 2021 for a similar consideration). This joint coordination, however, is at the expense of individual performance since participants reciprocally ceded timing, yielding tempo contagion. Second, emergent synchronization is driven by the same visuomotor information but the higher the cerebellar activity the more tempo contagion is prevented. It is known that the cerebellum has a prominent role in maintaining stable rhythms (Andersen & Dalal, 2021; Molinari et al., 2007), and specifically, its lateral portion integrates visual and motor information (for a review, see Tzvi, Loens, & Donchin, 2022). Thus, the cerebellum plays a crucial role in managing the temporal information stemming from visuomotor sources by interacting with high-level visual areas.

Functional connectivity analysis revealed reciprocal communication between the cerebellar and visual activity, suggesting that the cerebellum has a role in processing biological motion information (for instance, see, Sokolov et al., 2012). Our analysis confirmed an interplay between visual and motor information at least in the NI task: the higher the tempo contagion, the less the connectivity between the right cerebellum and a portion of the right LOTc or, said otherwise, the stronger the connectivity, the less the contagion (**Fig.5**). This finding indicates that individual cerebellar activity favors prevention to tempo contagion during visuomotor synchronization. Moreover, the analysis also showed a negative association with the connectivity between the cerebellum and a right fronto- parietal cluster (precentral/post-central gyri), compatible with the idea that entrainment occurs more easily the less cortical sensorimotor control. Both findings are in accord with previous evidence on the role of the cerebellum in visuomotor adaptation (Tzvi, Loens, & Donchin, 2022; Tzvi et al., 2020) and action observation (Sokolov et al., 2010). Last, in the JA task a plausible expectation was to observe connectivity between the LOTc and motor or premotor regions as an index of active co-regulation, but no connectivity pattern was associated with tempo contagion. This lack of information did not allow us to describe better tempo contagion in JA, nevertheless, it does not affect the interpretation of our results.

**Figure 5.**
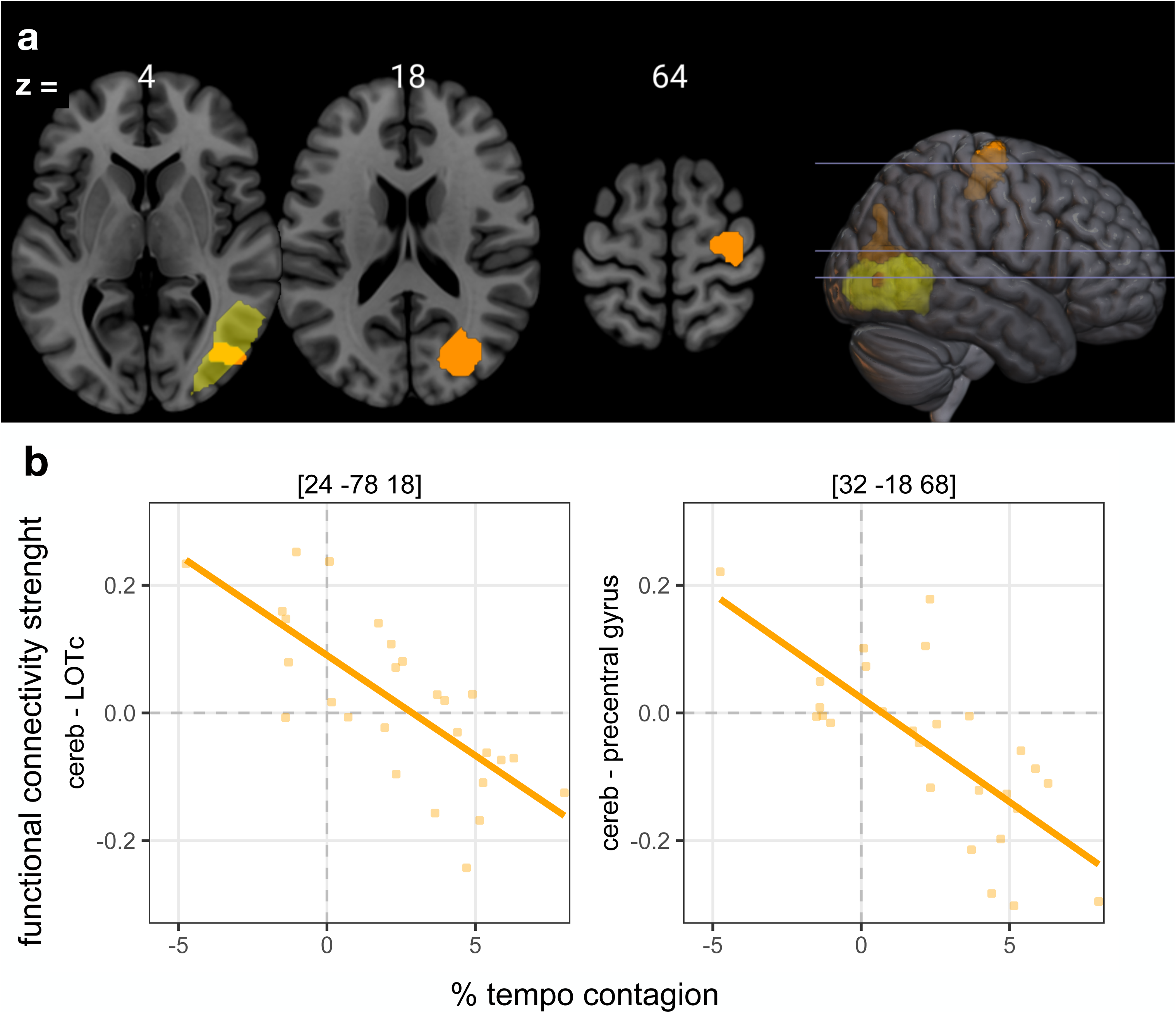
Results of the functional connectivity with the percent tempo contagion in the Non-Interactive condition. **a)** Orange clusters indicate the brain regions functionally connected with the right cerebellum cluster shown in the activation task (see **Fig.4a**). Note that the cluster in the lateral occipital cortex (LOTc, orange) overlaps with the cluster found in the activation task (yellow). **b)** Scatterplots of the negative linear relation between the functional connectivity strength and the % tempo contagion effects for both clusters. No significant results were found when correlating % tempo contagion effects from the Joint Action condition.

As a last remark, the neural network included also motor, premotor, and sensorimotor clusters known to be active during listening to rhythms (Bengtsson et al., 2009), comprising the inferior frontal gyrus, which is involved in action observation and imitation, especially for goal-directed actions (Molnar-Szakacs et al., 2005). Despite their established role in the motor control of rhythmic movements, our analyses did not show contributions from these areas to generate tempo contagion (see Fig.S2 in the supplemental online). At least for our findings, we suggest that visuomotor temporal adjustments occur automatically as sustained by perceptual-action matching mechanisms involving cerebellar and visuomotor activations and escaping the control exerted by the higher levels of the motor hierarchy implicated in action goal representations.

## Conclusion

Our work unveiled information on the neural correlates of emergent and planned interpersonal synchronization during a visuomotor task. On the one hand, extrastriate visual areas (e.g., the LOTc) are crucial for achieving joint synchronization. On the other hand, the cerebellum is crucial for maintaining an individual’s target timing while preventing entrainment to the partner’s movements. Despite the role of motor contagion being established in social interactions (for instance, see Blakemore & Frith, 2005), such a rivalry between emergent and planned processes is of likewise importance for the understanding of temporal contagion effects. These findings suggest a two-fold antagonist mechanism for preserving individual synchronization or promoting joint one, which may constitute a temporal requirement for self-other distinction (Decety & Sommerville, 2003) during motor interaction.

## Credit author statement

S.U.: conceptualization, methodology, investigation, data curation, formal analysis, visualization, writing - original draft, writing - review & editing. L.M.S.: conceptualization, methodology, formal analysis, writing original draft, writing - review & editing. C.DB.: investigation, data curation, formal analysis. F.D.: formal analysis, writing - review & editing. GB: writing - review & editing. E.P.: writing - original draft, writing - review & editing.

## Acknowledgements

We thank the technical staff of the San Gerardo Monza’s Hospital for making this study possible, in particular the radiologist technician Paola Lanzoni, and the Tecnomed Foundation (https://fondazionetecnomed.it/) that founded the magnetic resonance.

## Data availability

Data and supplemental materials are available at: https://osf.io/jvwz3/.

