## supplemental materials for "Neural mechanisms for visuomotor co-regulation in social synchronization"

**To synch or not to synch: neural correlates of tempo contagion during visuomotor interpersonal synchronization**

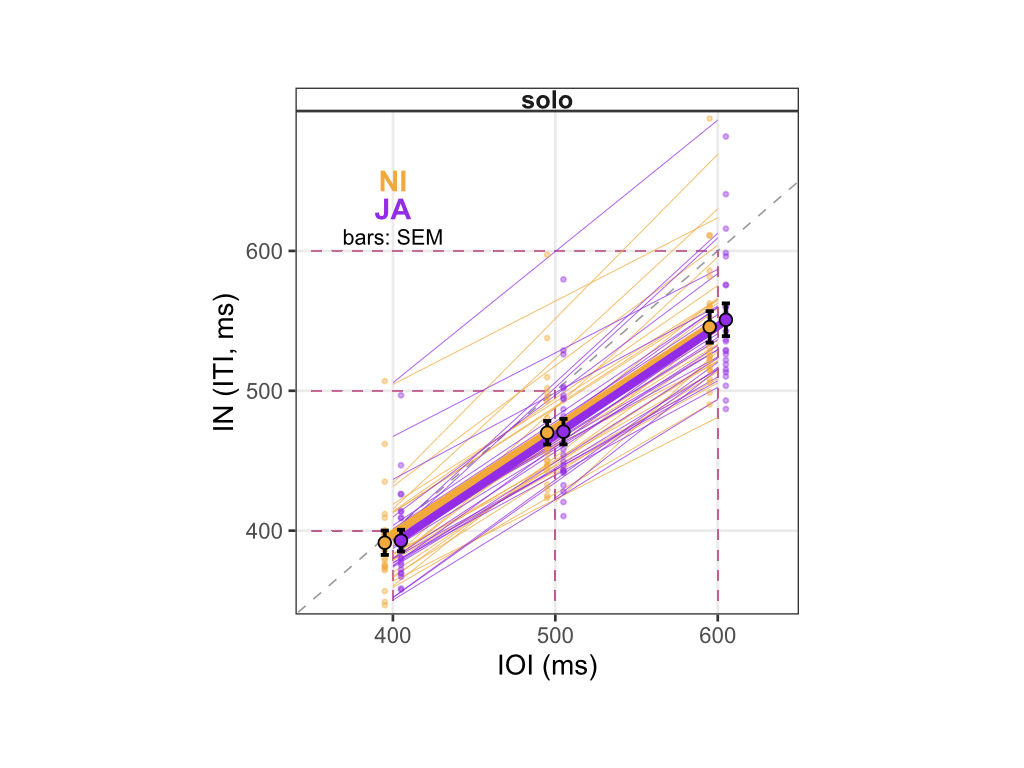

**Figure S1: Inter-tap-interval (ITI) in Solo trials.** Individual (small points) and group averages (large points) of IN participants are plotted as a function of the target inter-onset-interval (IOI). Colors identifies Solo trials of the Non-interactive (‘NI’, orange) and Joint Action (‘JA’, purple) fMRI runs. Bars are standard error of the mean (SEM). The plot shows that ITIs were longer for longer IOIs but the overall performance did not differ between runs.

Repeated-measures ANOVA (function *ezANOVA* from the *ez* package in R)

Dependent variable tested = difference between IN and OUT participants

IOI (IN participant) - IOI (OUT participant)

| **Predictor** | **DFn** | **DFd** | **F** | **P value** | **Generalized eta^2** |
| --- | --- | --- | --- | --- | --- |
| IOI (numeric) | 1 | 29 | 2.82 | 0.10 | 0.0162 |
| Condition (NI/JA) | 1 | 29 | 0.003 | 0.95 | 0.00072 |
| IOI * Condition | 1 | 29 | 2.59 | 0.11 | 0.0126 |

| **Brain area**  **(Brodmann)** | **Left hemisphere** |  |  |  | **Right hemisphere** |  |  |  |
| --- | --- | --- | --- | --- | --- | --- | --- | --- |
|  | **X** | **Y** | **Z** | **Z-score** | **X** | **Y** | **Z** | **Z-score** |
| **Right parietal cluster**  k = 693, p_FWE-corr_ = .020 |  |  |  |  |  |  |  |  |
| Inferior parietal gyrus (40) | -- | -- | -- | -- | 54 | -38 | 52 | 6.19* |
| **Fronto-parietal-insular and subcortical cluster**  k = 14362, p_FWE-corr_ < .001 |  |  |  |  |  |  |  |  |
| Postcentral gyrus | -46 | -14 | 58 | >8* | -- | -- | -- | -- |
|  | -38 | -20 | 54 | >8* | -- | -- | -- | -- |
| Precentral gyrus (4) | -36 | -24 | 56 | >8* | -- | -- | -- | -- |
| Precentral gyrus (6) | -34 | -24 | 68 | >8* | -- | -- | -- | -- |
|  | -56 | 6 | 18 | >8* | -- | -- | -- | -- |
|  | -58 | 2 | 38 | 7.81* | -- | -- | -- | -- |
|  | -60 | 6 | 30 | 7.73* | -- | -- | -- | -- |
| Supplemental motor area (6) | -4 | 0 | 60 | >8* | -- | -- | -- | -- |
| Supramarginal gyrus | -54 | -22 | 22 | >8* | -- | -- | -- | -- |
|  | -50 | -38 | 28 | 5.28* | -- | -- | -- | -- |
| Medial cingulate cortex | -- | -- | -- | -- | 12 | 14 | 34 | 4.26 |
| Insula | -42 | 2 | 4 | >8* | -- | -- | -- | -- |
| Pallidus | -22 | -2 | 0 | 5.17* | -- | -- | -- | -- |
| Thalamus | -16 | -18 | 6 | 4.36* | -- | -- | -- | -- |
| **Right fronto-insular cluster**  k = 1696, p_fwe-corr_ < .001 |  |  |  |  |  |  |  |  |
| Precentral gyrus (6) | -- | -- | -- | -- | 58 | 10 | 20 | 6.30* |
|  | -- | -- | -- | -- | 54 | 4 | 46 | 4.67* |
| Insula | -- | -- | -- | -- | 44 | 8 | 2 | 6.03* |
| **Right cerebellum cluster**  k = 2041, p_FWE-corr_ < .001 |  |  |  |  |  |  |  |  |
| Cerebellum, VI | -- | -- | -- | -- | 22 | -56 | -22 | >8* |
| **Right cerebellum cluster**  k = 673, p_FWE-corr_ < .001 |  |  |  |  |  |  |  |  |
| Cerebellum, VIII | -- | -- | -- | -- | 16 | -68 | -52 | 7.36* |

**Table S1. Simple effect of ‘Solo’ condition.** X, Y, and Z are the stereotactic coordinates of the activations in the MNI space. All reported voxels (p < .001uncorr.) are included in clusters surviving the FWER correction at the cluster-level. A maximum of 16 coordinates (local maxima) per cluster have been reported, each placed at 4 mm apart, as reports in SPM12. (*) Z-scores statistically significant also after whole brain FWER correction (p < .05) at the voxel-level (FWER: family-wise error rate; MNI: Montreal Neurological Institute).

| **Brain area**  **(Brodmann)** | **Left emisphere** |  |  |  | **Right hemisphere** |  |  |  |
| --- | --- | --- | --- | --- | --- | --- | --- | --- |
|  | **X** | **Y** | **Z** | **Z-score** | **X** | **Y** | **Z** | **Z-score** |
| **Left occipital cluster**  k = 1452, p_FWE-corr_ = .001 |  |  |  |  |  |  |  |  |
| Medial occipital gyrus (37) | -44 | -70 | 4 | >8* | -- | -- | -- | -- |
| Inferior occipital gyrus (18) | -30 | -94 | -6 | >8* | -- | -- | -- | -- |
| Inferior occipital gyrus (19) | -38 | -88 | -8 | 7.41* | -- | -- | -- | -- |
| **Right temporo-occipital cluster**  k = 1502, p_FWE-corr_ = .001 |  |  |  |  |  |  |  |  |
| Medial temporal gyrus (37) | -- | -- | -- | -- | 46 | -64 | 4 | >8* |
| Inferior occipital gyrus (19) | -- | -- | -- | -- | 34 | -90 | -2 | >8* |
| **Right parietal cluster**  k = 1680, p_FWE-corr_ < .001 |  |  |  |  |  |  |  |  |
| Inferior parietal gyrus (40) | -- | -- | -- | -- | 56 | -52 | 40 | 6.94* |
|  | -- | -- | -- | -- | 32 | -48 | 48 | 5.95* |
|  | -- | -- | -- | -- | 48 | -50 | 52 | 5.87* |
| Supramarginal gyrus | -- | -- | -- | -- | 64 | -40 | 30 | 5.14* |
| **Fronto-parietal cluster**  k = 16447, p_FWE-corr_ < .001 |  |  |  |  |  |  |  |  |
| Postcentral gyrus | -46 | -14 | 58 | >8* | -- | -- | -- | -- |
|  | -38 | -20 | 54 | >8* | -- | -- | -- | -- |
| Precentral gyrus (6) | -48 | -4 | 52 | >8* | 50 | 4 | 50 | 7.29* |
|  | -46 | 0 | 52 | >8* | 46 | 4 | 40 | 6.87* |
| Precentral gyrus (44) | -- | -- | -- | -- | 50 | 8 | 30 | 6.47* |
| Inferior parietal gyrus (40) | -52 | -48 | 50 | 7.70* | -- | -- | -- | -- |
|  | -36 | -48 | 50 | 7.10* | -- | -- | -- | -- |
| Supplemental motor area (6) | -6 | 0 | 62 | 7.45* | 10 | 6 | 62 | 6.91* |
|  | -- | -- | -- | -- | 12 | 18 | 56 | 6.56* |
|  | -- | -- | -- | -- | 16 | 14 | 64 | 6.56* |
| Supramarginal gyrus | -48 | -40 | 28 | 6.77* | -- | -- | -- | -- |
|  | -52 | -24 | 24 | 6.31* | -- | -- | -- | -- |
| Inferior frontal gyrus, pars opercolaris (6) | -56 | 8 | 18 | 5.70* | -- | -- | -- | -- |
| **Left subcortical cluster**  k = 531, p_FWE-corr_ = .044 |  |  |  |  |  |  |  |  |
| Pallidus | -20 | 0 | 2 | 5.27* | -- | -- | -- | -- |
| Thalamus | -14 | -18 | 4 | 4.12 | -- | -- | -- | -- |
| **Left cerebellum cluster**  k = 634, p_FWE-corr_ = .026 |  |  |  |  |  |  |  |  |
| Cerebellum, Crus 1 | -34 | -64 | -30 | 5.76* | -- | -- | -- | -- |
|  | -40 | -60 | -32 | 5.55* | -- | -- | -- | -- |
| **Right cerebellum cluster**  k = 1500, p_FWE-corr_ = .001 |  |  |  |  |  |  |  |  |
| Cerebellum, VI | -- | -- | -- | -- | 20 | -54 | -24 | 7.03* |
| Cerebellum, VI | -- | -- | -- | -- | 26 | -58 | -26 | 6.91* |
| **Right cerebellum cluster**  k = 650, p_FWE-corr_ = .025 |  |  |  |  |  |  |  |  |
| Cerebellum, VIIb | -- | -- | -- | -- | 10 | -74 | -48 | 5.75* |
| Cerebellum, VIII | -- | -- | -- | -- | 20 | -70 | -54 | 5.30* |
|  | -- | -- | -- | -- | 24 | -68 | -54 | 5.30* |

**Table S2. Effect of the ‘Together > Solo’ linear contrast.** X, Y, and Z are the stereotactic coordinates of the activations in the MNI space. All reported voxels (p < .001uncorr) are included in clusters surviving the FWER correction at the cluster-level. A maximum of 16 coordinates (local maxima) per cluster have been reported, each placed at 4 mm apart, as reports in SPM12. (*) Z-scores statistically significant also after whole brain FWER correction (p < .05) at the voxel-level (FWER: family-wise error rate; MNI: Montreal Neurological Institute).

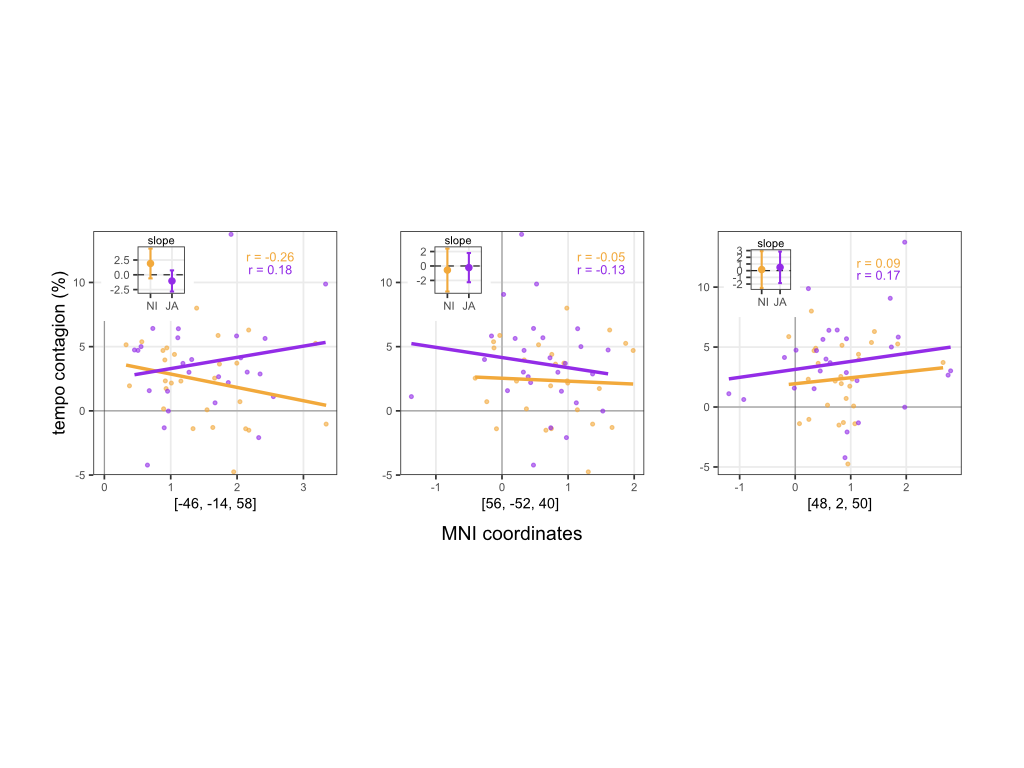

**Figure S2.** Linear regression between the individual % tempo contagion (y-axis) and the BOLD signal (x-axis) extracted at the local maxima MNI coordinates (left plot: left frontoparietal cluster; middle: right parietal cluster; right plots: frontal right cluster; see Fig.4a in the main text). Each point represents a different IN participant. Inset plots: 95% confidence intervals (CIs) around slopes extracted from the linear mixed model providing the best fit (see text for de tails). Results reveal that these brain regions did not modulate % tempo contagion (see details about modelling below). Orange: Non-interactive (NI); purple: Joint Action (JA). Data is plotted by applying the same threshold reported in Table 1 (puncorr. < .001 at the voxel level and pFWER-corr. < .05 at the cluster-level; see main text).

**ANCOVA model: % tempo contagion ~ BOLD * Condition (NI/JA)**

*Left fronto-parietal cluster*

| Predictor | Estimate | Std.Error | t value | P-value |
| --- | --- | --- | --- | --- |
| Intercept (% tempo contagion) | 3.9 | 1.47 | 2.65 | 0.01 |
| Main effect BOLD | -1.03 | 0.88 | -1.17 | 0.24 |
| Main effect Condition | -1.48 | 2.06 | -0.72 | 0.47 |
| BOLD * Condition interaction | 1.91 | 1.25 | 1.5 | 0.13 |

*Right parietal cluster*

| Predictor | Estimate | Std.Error | t value | P-value |
| --- | --- | --- | --- | --- |
| Intercept (% tempo contagion) | 2.5 | 1.01 | 2.5 | 0.015 |
| Main effect BOLD | -0.22 | 1 | -0.22 | 0.82 |
| Main effect Condition | 1.61 | 1.36 | 1.18 | 0.24 |
| BOLD * Condition interaction | -0.56 | 1.48 | -0.38 | 0.7 |

*Right frontal cluster*

| Predictor | Estimate | Std.Error | t value | P-value |
| --- | --- | --- | --- | --- |
| Intercept (% tempo contagion) | 1.94 | 1.19 | 1.6 | 0.11 |
| Main effect BOLD | 0.49 | 1.17 | 0.42 | 0.67 |
| Main effect Condition | 1.19 | 1.49 | 0.8 | 0.42 |
| BOLD * Condition interaction | 0.16 | 1.36 | 0.12 | 0.9 |

**Functional connectivity analysis: Additional information**

*Preprocessing and denoising*

Data analyses were performed in MATLAB R2019b (Mathworks. Natick. MA. USA) using the software CONN^11^ and SPM12.Functional data were realigned using SPM realign & unwarp procedure, where all scans were coregistered to a reference image (first scan of the run) using a least squares approach and a 6 parameter (rigid body) transformation and resampled using b-spline interpolation to correct for motion and magnetic susceptibility interactions. Temporal misalignment between different slices of the functional data was corrected following SPM slice-timing correction procedure, using sinc temporal interpolation to resample each slice BOLD timeseries to a common mid-acquisition time.

Potential outlier scans were identified using ART as acquisitions with framewise displacement above 1 mm or global BOLD signal changes above 3 standard deviations. On average, excluded volumes were 2.14%. Only one participant who showed 135 outliers (31.1%) was excluded from the sample. Functional data were normalized using an indirect normalization procedure. Functional images were coregistered with a T1-weighted structural image of each participant, which was then segmented and stereotactically normalized into the SMP12 template to allow for group analyses. Deformation fields used for T1 segmentation were then applied to coregistered functional scans; here, the data matrix was interpolated to produce voxels 2x2x2 mm in dimension. Last, the stereotactically normalized scans were smoothed using a Gaussian filter of 10 x 10 x 10 mm to improve the signal-to-noise ratio, making the data suited for cluster-level correction for multiple comparisons.

In addition, functional data were denoised by including the regression of potential confounding effects characterized by white matter timeseries (5 CompCor noise components), CSF timeseries (5 CompCor noise components), motion parameters and their first order derivatives (12 factors), outlier scans, task activation effects and their first order derivatives, and linear trends, followed by high-pass frequency filtering of the BOLD timeseries above 0.008 Hz. CompCor noise components within white matter and CSF were estimated by computing the average BOLD signal as well as the largest principal components orthogonal to the BOLD average, motion parameters, and outlier scans within each subject's eroded segmentation masks.
